# Examining perfluorohexane sulfonate (PFHxS) impacts on sensorimotor and circadian rhythm development

**DOI:** 10.1101/2024.10.08.617320

**Authors:** Syed Rubaiyat Ferdous, Alfredo Rojas, Cole Frank, Heidi M. Sabatini, Xixia Luo, Sunil Sharma, Ryan Thummel, Christopher Chouinard, Subham Dasgupta

## Abstract

Perfluorohexane sulfonate (PFHxS) is a ubiquitous perfluoroalkyl substance known for its environmental persistence and potential toxicity. This study investigated PFHxS’s impact on zebrafish embryos, focusing on sensorimotor behavior, circadian rhythm disruption, and underlying molecular mechanisms. Under 24 hr dark incubations, PFHxS exposure induced concentration-dependent hyperactivity within larval photomotor response, characterized by the distinctive “O-bend” response, strong light-phase hyperactive movement and seizure-like movements. It appears that PFHxS-treated embryos cannot sense light cues in a normal manner. Similar hyperactivity was seen for acoustic startle response assay, suggesting that the response is not merely visual, but sensorimotor. LC-MS studies confirmed detectable uptake of PFHxS into embryos. We then conducted mRNA-sequencing across multiple time points (48 and 120 hpf) and concentrations (0.00025, 0.0025 and 25 µM). Data at the 25 µM (2-120 hpf) exposure showed disrupted pathways associated with DNA and cell cycle. Interestingly, data at 0.00025 µM - an environmentally relevant concentration- at 48 hpf showed disruption of MAPK and other signaling pathways. Immunohistochemistry of eyes showed reduced retinal stem cell proliferation, consistent with observed DNA replication pathway disruptions. To assess if these impacts were driven by circadian rhythm development, we manipulated light/dark cycles during PFHxS incubation; this manipulation altered behavioral patterns, implicating circadian rhythm modulation as a target of PFHxS. Since circadian rhythm is modulated by the pineal gland, we ablated the gland using metronidazole; this ablation partially rescued hyperactivity, indicating the gland’s role in driving the phenotype. Collectively, these findings underscore proclivity of PFHxS to cause neurodevelopmental toxicity, necessitating further mechanistic exploration and environmental health assessments.

## Introduction

Perfluorohexane sulfonate (PFHxS), a perfluoroalkyl substance (PFAS), has emerged as a topic of profound concern within the field of environmental toxicology, driven by its persistent presence in the environment and its potential adverse effects on living organisms. PFAS are widely used in firefighting foams, carpets, papers, and textiles for their heat and water-resistant properties^1,2^. PFHxS’s strong carbon-fluorine bonds contribute to its thermal, chemical, and biological stability, leading to its widespread presence in environmental matrices and living organisms^3–5^. Although industries have shifted to alternative, lower molecular-weight compounds due to safety concerns^6^, PFHxS concentrations in surface and source waters range from 0.0005 µM to 0.15 µM, while concentrations in drinking water have been reported to reach up to 0.00039 µM^7^. High levels have been found in industrial or military areas, such as 0.33 µM in Oakey, Australia, posing health risks^8^. PFHxS is a prevalent environmental contaminant detected in human serum, breast milk, and various tissues and decreases serum thyroxine (T4) levels in rats^9^. PFHxS exposure is linked to liver disease, diabetes, and behavioral changes, with animal studies, including zebrafish research, showing it causes endocrine disruption, neurotoxicity, and developmental disorders^10–16^. For instance, Zebrafish exposed to 20 μM of PFHxS exhibited enlarged body length and yolk sac area, showing morphometric changes^12^. Conversely, neurotoxicity was observed fish exposed to 14 μM PFHxS^17^. Another study revealed that exposure to 10 μM of PFHxS was linked with alterations in lipids that stimulate oxidative stress, changes in beta-oxidation and inflammation^18^. These findings underscore the significant impact of PFHxS on health and the environment, specifically from a developmental perspective.

This study focuses on understanding the basis of unique sensorimotor and seizure-like hyperactivity patterns displayed by PFHxS-exposed zebrafish embryos and how these phenotypes are dependent on circadian rhythm changes. Circadian rhythm constitutes the body’s biological clock and is regulated by the eye and pineal gland^19^. Leveraging mRNA sequencing, immunohistochemistry, analytical chemistry and behavioral assays^20^, our study provides, for the first time, strong evidence of PFHxS-induced disruption of circadian rhythm and identifies gene-level disruptions at environmentally relevant concentrations.

## MATERIALS AND METHOD

### Zebrafish husbandry

Wild-type adult zebrafish (5D) were obtained from the Sinnhuber Aquatic Research Laboratory. They were maintained at 28 ± 2°C with a 14h/10h light/dark photoperiod on a recirculating system within the Aquatic Animal Research Laboratory at Clemson University. Breeding and exposures were conducted under Clemson Institutional Animal Care and Use Committee (IACUC) protocols (AUP2022-0434 and AUP2023-0114).

### Chemicals

PFHxS (97% purity) (CAS #: 79-94-7) and Metronidazole (CAS #: 443-48-1) were purchased from Sigma-Aldrich and stored in dimethyl sulfoxide (DMSO). The working solution was prepared in 1X embryo media (0.17 mM KCl, 0.33 mM CaCl_2_, 0.33 mM MgSO4 and 5 mM NaCl, pH 7). All chemicals and reagents used in the present study were of analytical grade.

### Larval photomotor response (LPR) and acoustic larval startle response (LSR)

Experiments were conducted in 96-well plates (Corning), with one embryo and 100 μl test solution per well and 32 embryos per treatment. Embryos were exposed to 0.00025, 0.0025, 0.025, 0.25, 2.5, 3.125, 6.25, 12.5 and 25 µM of PFHxS under 24 hr dark (24D), 24 hr light (24L), or 14 hr light:10h dark (14L:10D) incubation conditions for circadian rhythm variation, from 2 to 120 hpf. At 120 hpf, LPR and LSR assays were conducted using a ZebraBox (Viewpoint, France). Detailed description of assays is provided in the Supplemental file. Data were obtained from the Zebralab software and analyzed using a combination of R and GraphPad Prism 10.

### Assessments of seizure-like behavior

The seizure-like behavior of control or PFHxS-treated fish was analyzed from ZebraLab videos in a blinded manner using adopted methods^21,22^. Details are provided in the Supplemental file.

### Quantification of PFHxS uptake using LC-MS Analysis

PFHxS levels in 5 dpf zebrafish larvae exposed to 0, 0.0025 or 25 µM PFHxS from 2 hpf to 120 hpf were measured using a modified protocol from a prior publication^15^. We selected 0.00025 µM and 0.0025 µM to reflect environmentally relevant concentrations, while 25 µM was chosen to assess acute toxicity and mechanistic effects. Embryos were collected (150 embryos per replicate pool, 3 replicate pools) and processed for LC-MS using an Agilent 1290/6560 as described in Supplemental file. PFHxS concentration was determined by the extracted peak area relative to a calibration curve.

### mRNA sequencing

Embryos were exposed to 0, 0.00025, 0.0025, and 25 μM PFHxS in 96 well plates and incubated at 28 ºC from 2-48 or 2-120 hpf, similar to the setup for LPR assay. RNA was extracted (N= 4 replicate pools of 5 embryos per pool) using a Direct**-**zol**™** RNA MiniPrep Kit. Library preparation and sequencing were performed by Novogene (California, USA). Gene Ontology (GO) enrichment analyses were performed on differentially expressed genes using iDEP (http://bioinformatics.sdstate.edu/idep96/). An FDR cutoff of 0.05 and a minimum fold change of 1.5 were applied to determine significance. Extended details as are provided in Supplemental File.

### Immunohistochemistry

Immunohistochemistry was performed on 16-micron retinal sections from treated 5 dpf larvae and controls using antibodies against Zpr-1, Rhodopsin, and PCNA. TUNEL analysis was conducted to assess apoptosis. Detailed methods are provided in the Supplemental File.

### Pineal gland ablation experiments

Metronidazole (Mtz) is a nitroimidazole that can ablate pineal gland development in zebrafish when exposed between 24 and 48 hpf ^23^. To study the impacts of pineal gland ablation, embryos were co-exposed to 25 μM PFHxS (2-120 hpf) and 20 mM Mtz (24-48 hpf) as described in the Supplemental file. At 120 hpf, LPR and LSR were conducted as described previously.

### Statistical analysis

All statistical analyses were conducted, and graphs were generated using GraphPad Prism 10.0. For normalized data, one-way analysis of variance (ANOVA) followed by Dunnett’s posthoc test p<0.05). An unpaired t-test was used where there was not more than one treatment group (DMSO vs 25 µM, PFHxS). For metronidazole data, statistical analyses were conducted using 2-way ANOVA, followed by Tukey’s posthoc test, p<0.05. For seizure assay, statistically significant was determined using one-way ANOVA, followed by Dunnett’s test, with p < 0.05 considered significant.

## Results and Discussion

The circadian rhythm and visual system development are intertwined, and disruption of one can affect the other^24^. PFHxS is a ubiquitous PFAS, and prior studies have shown that it impacts visuomotor behavior^17^ but we delve into the characteristics of swimming patterns and how they are impacted by retinal development and circadian rhythm. PFHxS-exposed zebrafish embryos raised in 24 hr darkness exhibited significant hyperactivity during the light epoch and a hyperactive “O-bend” response during the LPR assay (**Figure 1A**), and this response was concentration-dependent in both epochs (**Figure 1B, 1C**), although the dark epoch showed a U-shaped response. Based on these phenotypes, it appears that the fish cannot sense light. Normally, zebrafish larvae exhibit a clear light-dark response, where they become more active during dark periods and significantly reduce their movement or ‘freeze’ when exposed to light. This behavior is a well-established survival mechanism in zebrafish, as the sudden exposure to light typically signals potential threats. In our experiments, PFHxS-exposed larvae did not exhibit the expected reduction in activity during light phases, suggesting an impaired ability to detect or process light cues, further supporting the hypothesis of sensory disruption. When exposures were initiated at differing days post fertilization, hyperactivity was seen in all groups except 96 – 120 hpf **(Figure 1D, 1E**), suggesting that even a 2-day exposure is sufficient to elicit this response. The acoustic startle response (LSR) mimicked light epoch LPR, with concentration-dependent hyperactivity (**Figure 1F**) and hyperactivity on exposure initiation up to 72 hpf (**Figure 1G**). Exposure to anxiogenic compounds is typically associated with an augmentation of swimming activity during dark periods, potentially attributed to interference with sensory or dopaminergic systems^25^. Exposure to PFAS has been documented to induce behavioral toxicity in animal studies^26–30^ with LPR hyperactivity^26,31^ and diminished habituation^27^. Specifically, hyperactivity stemming from PFHxS exposure was also reported by prior work^17,32^. Our study is consistent with these responses but critically parses out the phasic swimming behavior (O-bend versus regular swimming) and demonstrates that PFHxS exposure under 24 hr dark introduces consistent O-bend, not seen in 24 hr dark controls. These responses are not merely visuomotor, but also sensorimotor, as demonstrated by LSR mimicking LPR. PFHxS-exposed embryos in our study also displayed seizure-like movements (**Figure 1H**), suggesting that neurotransmitters such as GABA may be a target^33^. LC-MS confirmed significant PFHxS uptake at 25 µM and 0.0025 µM-an environmentally low concentration **(Figure 1I)**, but surprisingly, the PFHxS difference between two nominal concentrations 10,000X apart was only ∼21X. This suggests that even at a high 25 µM concentration, PFHxS uptake is limited, perhaps due to chorion presence during the first 2 days.). In our RNA-seq analysis, 27 genes were differentially expressed at 0.0025 µM (Figure 2A), compared to 2,462 genes at 25 µM (Figure 2B). Importantly, concentrations of 0.00025 and 0.0025 µM used in the experiment reflect environmentally relevant levels of PFHxS found in aquatic environments, ranging from 0.0005 µM to 0.15 µM, with higher concentrations observed in specific industrial or military sites^7,8,34^.

**Figure 1.**
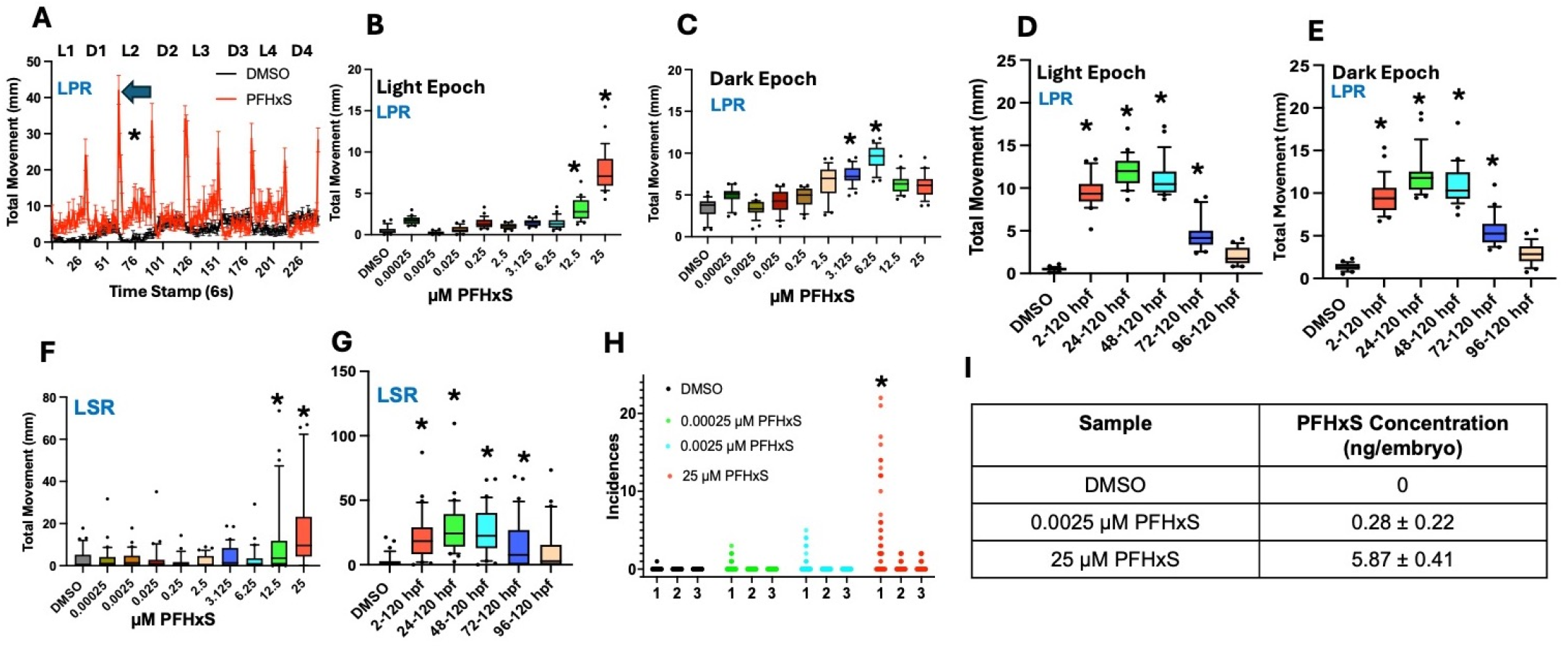
PFHxS exposure induces hyperactivity and seizure-like behavior in zebrafish embryos and larvae. **(A)** Embryos raised in 24 hr darkness exhibited hyperactivity in the light epoch, and hyperactive “O-bend” (Blue arrow). Embryos were exposed to 25 µM PFHxS from 2 to 120 hpf. L1-L4 indicate 4 light epochs, D1-D4 indicate 4 dark epochs. **(B, C)** Concentration dependence of PFHxS-induced hyperactivity. Embryos were exposed to 0 to 25 µM PFHxS. **(D, E)** Time-dependent hyperactivity in zebrafish larvae following PFHxS exposure. Embryos/larvae were exposed to 25 µM PFHxS at different life stages, starting from 2 hpf up to 96 hpf. **(F, G)** LSR following acoustic stimulus elicits hyperactivity in PFHxS-treated Zebrafish. Embryos/larvae were exposed to 25 µM PFHxS at different time points, starting from 2 hpf up to 96 hpf, and exposed to a 600 Hz sound for 900 ms. **(H)** Embryos exposed to PFHxS in 24 hr dark exhibited dose-dependent seizure-like movements. Three stages of behavior: hyperlocomotion, whirlpool swimming, and tonic seizure with brief convulsions and loss of posture followed by immobility. **(I)** LC-MS Analysis confirmed uptake of PFHxS in zebrafish embryos. * statistically significant based on one way ANOVA, followed by Dunnett’s test (p<0.05), N=32.

**Figure 2.**
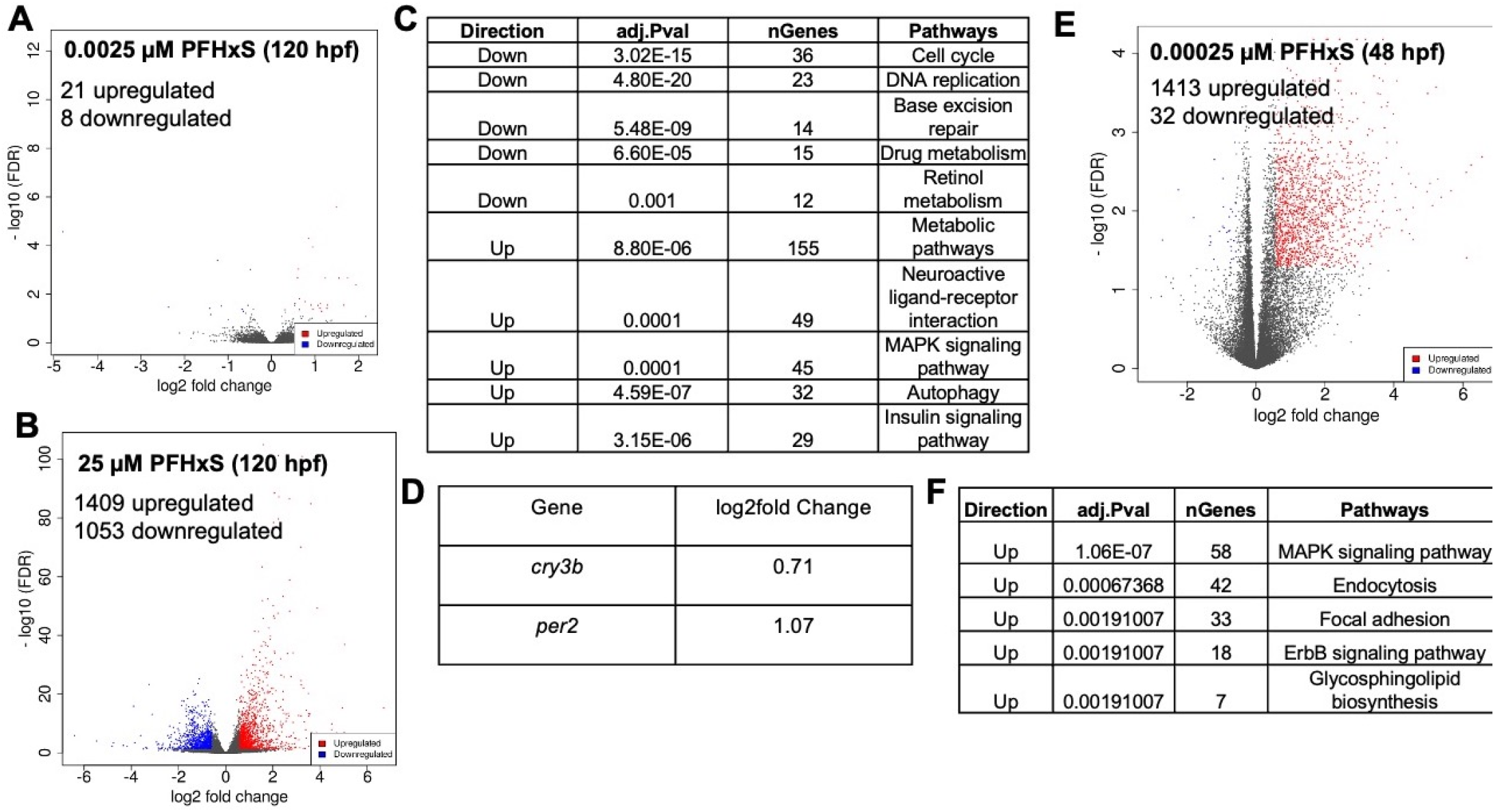
mRNA-sequencing and gene ontology analyses in embryos exposed to 0, 0.0025 or 25 µM and 0.00025 µM PFHxS from 2 to 48 or 120 hpf in 24 hr dark incubations. **(A, B)** volcano plot for differentially expressed gene from embryos exposed to 0, 0.0025 or 25 µM PFHxS from 2 to 120 hpf. **(B)** KEGG pathway analysis indicated various affected pathways from 120 hpf embryos exposed to 0, 0.0025 or 25 µM PFHxS. **(D)** Genes related to circadian rhythm exhibit differential expression in 25 µM PFHxS exposed 120 hpf embryos. **(E)** Volcano plot for differentially expressed gene from 48 hpf embryos exposed to 0.00025 µM PFHxS. **(F)** KEGG pathway analysis indicated various affected pathways from 48 hpf embryos exposed to 0.00025 µM PFHxS. An FDR cutoff of 0.05 and a minimum fold change of 1.5 were applied to determine significance.

Based on our RNA-seq study, KEGG pathway analysis at 25 µM **(Figure 2C)** revealed decreased mRNA levels in DNA replication and cell cycle pathways, suggesting that PFHxS may impact cell division. In addition, the retinol metabolism pathway was downregulated; retinol is essential for synthesizing rhodopsin-the pigment in the retina crucial for low-light and color vision, indicating a potential link between PFHxS exposure and disrupted visual development. Furthermore, circadian rhythm mRNA levels (*cry3b, per2*) were increased **(Figure 2D)**. Both ***cry3b*** and ***per2*** are key components of the molecular circadian clock. ***cry3b*** (cryptochrome 3b) regulates circadian rhythms by inhibiting the activity of clock genes during the light phase, contributing to the maintenance of daily behavioral and physiological cycles. ***per2*** (period 2) plays a crucial role in the negative feedback loop of the circadian clock, regulating the timing and amplitude of circadian rhythms. Changes in the expression of these genes can disrupt normal circadian rhythms, leading to alterations in light-dependent behaviors and biological processes^19^. A separate transcriptomic analysis of 0.00025 µM-exposed embryos (2-48 hpf) revealed a different set of differentially expressed genes **(Figure 2E)** with pathways related to MAPK signaling, endocytosis, focal adhesion, ErbB signaling, and glycosphingolipid biosynthesis, suggesting an impact on protein synthesis and stress responses during earlier stages **(Figure 2F)**. Full datasets for DEGs and KEGG analysis results are provided in Supplemental Tables. These results highlight the significant impact of PFHxS on essential cellular processes and stress response mechanisms. Importantly, extensive gene-level disruptions are observed under environmentally relevant concentrations at which we do not see phenotypes, warranting further investigation into their implications. Collectively, these necessitate further investigation into mechanisms of PFHxS toxicity, such as cell division, coupled with examining the long-term health effects of developmental transcriptomic disruptions.

Based on phenotype and sequencing data, we investigated the impacts on visual development and circadian rhythm. Immunohistochemistry of the eye showed that PFHxS exposure resulted in a fewer number of proliferating cells in the ciliary marginal zone of the retina **(Figure 3A)**, a stem cell niche that gives rise to all retinal cell types except rod photoreceptors. This is consistent with PFHxS impacts on DNA replication and cell cycle pathways. Rods and cone cells are not well developed at these stages and were not impacted **(Figure S1)**. Variation of circadian rhythm by altering light/dark photoperiods showed that light epoch hyperactivity (observed in 24D) was mitigated under 14L:10D (**Figure 3B**). No statistically significant differences in activity were observed in 24L (**Figure 3B**). Collectively, these potentiate circadian rhythm as target of PFHxS. Notably, control embryos in 24L and 24L:14D, but not 24D, displayed O-bends **(Figure S2)**, suggesting that the presence of O-bends in the 24D PFHxS is a circadian rhythm anomaly that needs to be interrogated. To further assess the impact of the pineal gland (a regulator of circadian rhythm through melatonin production), Mtz co-exposure was used for pineal ablation; this partially rescued both LPR light epoch hyperactivity **(Figure 3C)** and LSR hyperactivity **(Figure 3D)**, suggesting a contributory role of pineal gland in these behaviors and potential for dose-dependent therapeutic interventions using Mtz. Further research is needed to elucidate the precise mechanisms by which the pineal gland influences these behaviors and to explore optimal intervention strategies.

**Figure 3.**
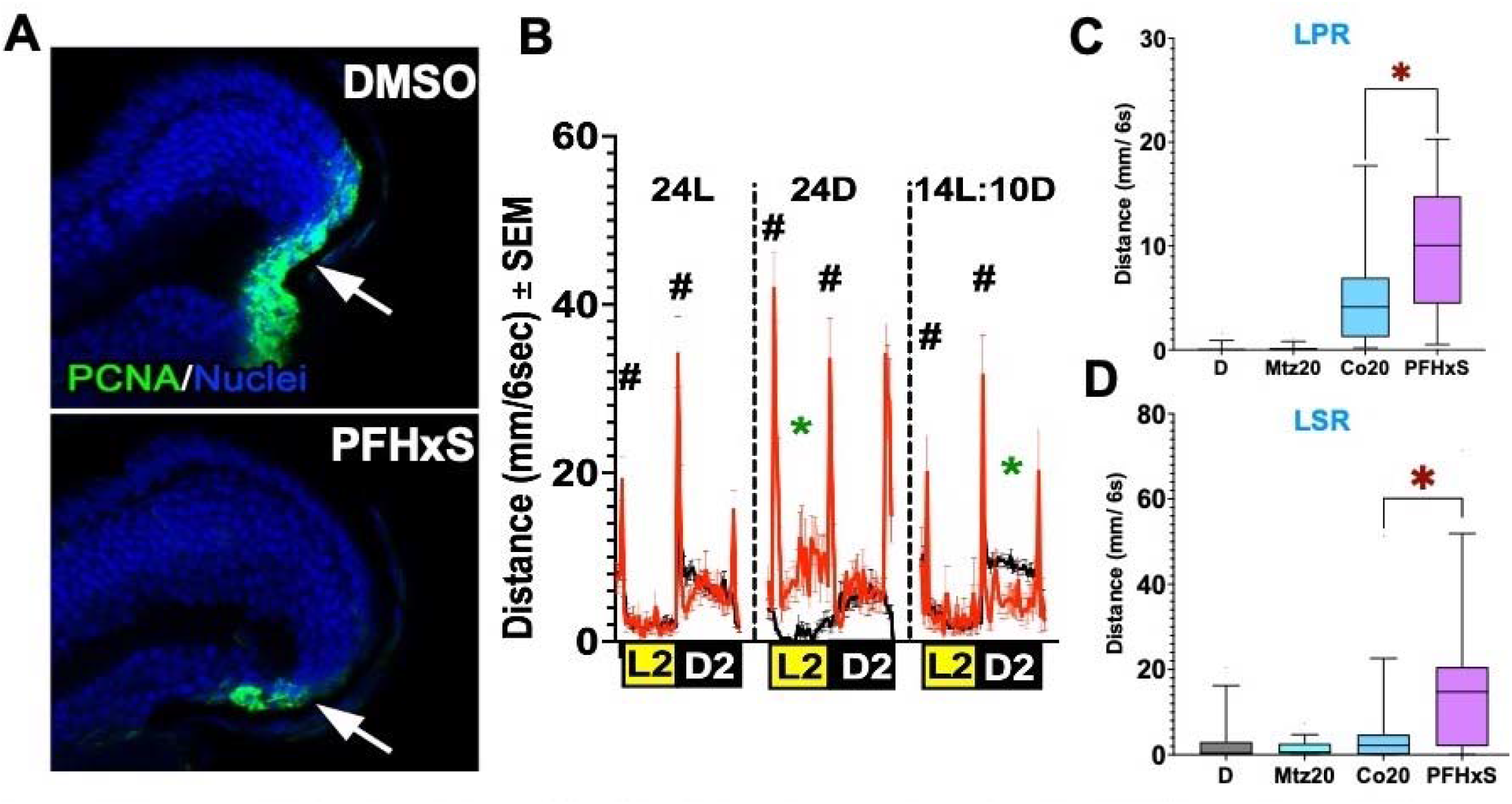
Immunohistochemistry and metronidazole experiments. **(A)** PCNA-positive cells in the ciliary marginal zone of the retina (white arrows) show a reduced number of proliferating cells in the 25 µM PFHxS-treated retinas. **(B)** LPR profiles under varied light/dark conditions. 24D-24 h dark, 14L: 10D-14 hr light and 10 hr dark and 24L-24 hr light incubations. These are distinct from L2 and D2, which represent the 2nd light and dark epochs of LPR assay. * statistically different movement based on t-test p<0.05. ^#^indicates significance for transition O-bends. The black line represents DMSO, and the red line represents 25 µM PFHxS. N=32. **(C, D)** Metronidazole co-exposure studies; D-DMSO; Mtz20-20 mM Metronidazole; PFHxS-25 µM PFHxS; Co20-Mtz-PFHxS coexposure. (C) LPR during the 2nd light epoch and (D) LSR. *statistically significant based on two-way ANOVA, followed by Tukey’s posthoc test p<0.05. N=32.

### Implications

Using a systemic approach, this study reveals that the effect of PFHxS exposure on sensorimotor development depends on circadian rhythm. These findings underscore how PFHxS exposure during development can interfere with visual and sensorimotor functions, potentially serving as precursors for long-term neurological health effects. While our data show transcriptomic changes at low PFHxS concentrations, it is important to note that these effects may be adaptive responses rather than predictors of long-term health outcomes, as no direct evidence of long-term impacts is demonstrated in our study. Understanding the long-term consequences of PFHxS exposure would require longitudinal studies that track developmental, behavioral, and physiological changes over the lifespan of exposed organisms. A possible experimental design could involve long-term exposure studies with zebrafish or other model organisms, followed by multi-generational assessments of neurodevelopment, reproductive health, and epigenetic changes to determine if early-life transcriptomic alterations translate into adverse health outcomes. Since circadian rhythm aberrations can impact overall health, understanding the long-term consequences of PFHxS exposure is crucial for developing regulatory measures to mitigate its adverse impacts on health and the environment.

## Acknowledgements

This research was made possible with startup funds to SDG and CC from Clemson University, the Clemson University SC STEM initiative to CC, Kids Without Cancer Foundation and the Children’s Foundation grants to RT and NIH/NIEHS grants R03-ES036327 and R03-ES036327-01S1 to SDG. We acknowledge Mr. John Smink, Facility Manager, Aquatic Animal Research Laboratory for husbandry of zebrafish and Dr. Robyn Tanguay (Oregon State University) for generously providing us with 5D founder fish. We also acknowledge Clemson University Genomics and Bioinformatics Facility for using a Tape station and Nanodrop (NIH grant numbers P20GM146584 and PM20GM139769).

## Supporting information

Extended experimental details of LPR and LSR assays, seizure-like behavior assessments, UPHLC-MS methods, RNA-seq, IHC and pineal gland ablation, as well as extended figures containing retinal IHC and circadian rhythm controls are presented within supplemental file. RNA-seq read counts, differentially expressed genes and KEGG pathway analyses from sequencing data are presented within supplemental tables.

### AI use

Chat GPT was used to paraphrase and furnish texts.

## Notes

### Competing Interest Statement

The authors have declared no competing interest.

