## Supplemental File for "Examining perfluorohexane sulfonate (PFHxS) impacts on sensorimotor and circadian rhythm development"

Number of Figures- 2

Number of Tables in this file- 3

Number of Sheets in Supplemental Table- 7

**Legends for Supplemental Table:**

Table S1: Read counts for 120 hpf data. Low- 0.0025 µM PFHxS, High- 25 µM PFHxS

Table S2: Differentially expressed genes for High treatment.

Table S3: KEGG pathway analysis for High treatment

Table S4: Differentially expressed genes for Low treatment.

Table S5: Read counts for 48 hpf data. PFHxS indicates 0.00025 µM PFHxS.

Table S6: Differentially expressed genes at 48 hpf.

Table S7: KEGG pathway analysis at 48 hpf.

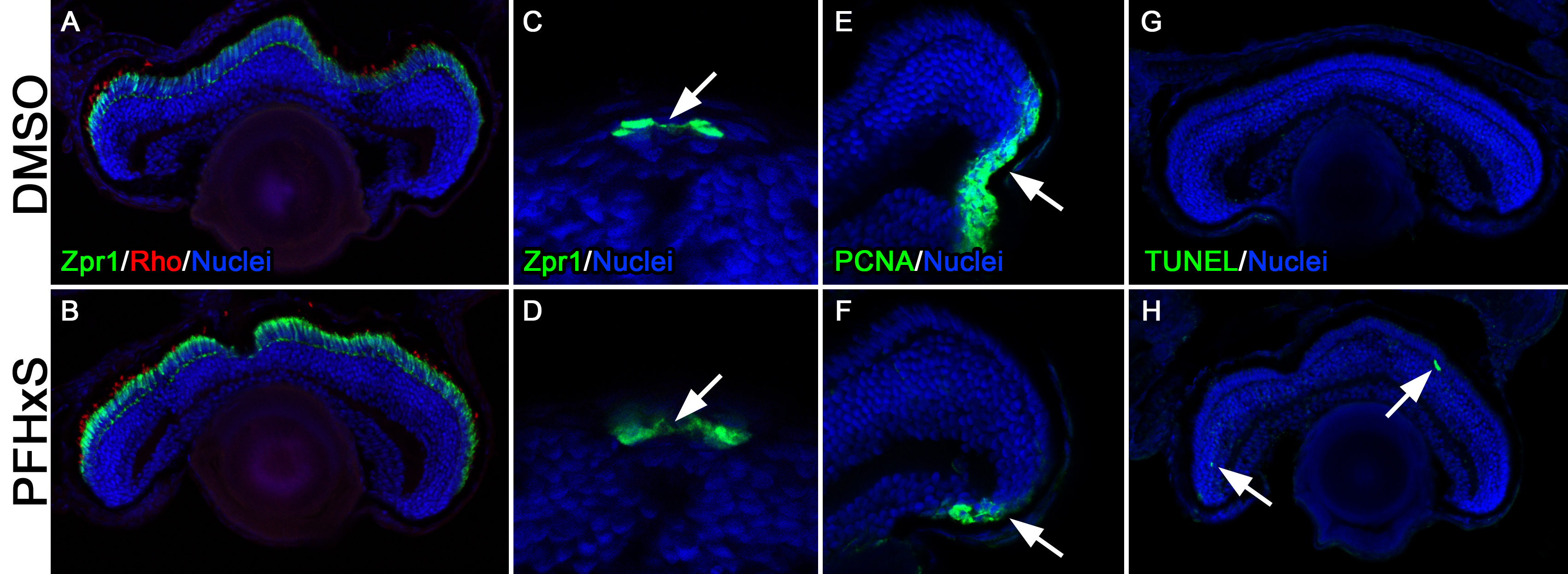

**Figure S1**. Immunohistochemical analysis of DMSO and PFHxS-treated larval at 5 dpf. (A-B) Retina section stained with Zpr1 (Arrestin 3; cones) and Rhodopsin (Rho; rods) shows normal formation of cone and rod photoreceptors in PFHxS-treated retinas. (C-D) Pineal sections in the forebrain stained with Zpr1 shows normal pineal formation in both groups. (E-F) PCNA-positive cells in the ciliary marginal zone of the retina show a reduced number of proliferating cells in the PFHxS-treated retinas. (G-H) TUNEL labeling of apoptotic cells shows minimal cell death in both treatment groups.

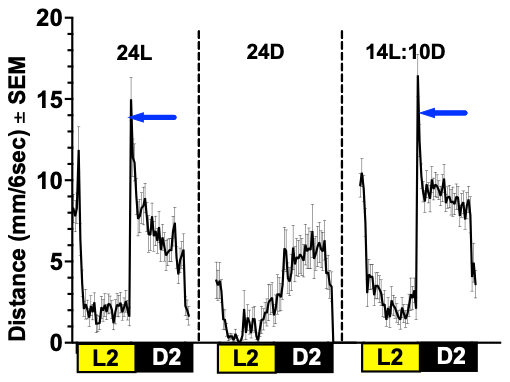

**Figure S2. Control LPR curves under different incubations. Blue arrows indicate O-bends, present in 24L and 14L:10D, but not in 24D.**

**1. Extended methods- LPR (Larval photo-motor response) and LSR (Larval startle response)**

**Experiments were conducted in a 96-well plate where each well had 100** µ**L of solution and one embryo.** Throughout the exposure phase, the 96-well plate housing zebrafish embryos remained in an incubator, sustaining a temperature of 28 ºC. This controlled setup aimed to replicate conditions suitable for studying the impact of PFHxS on zebrafish larvae during a critical developmental window. At 120 hpf embryos larval photo motor response was assessed on a Zebrabox (Viewpoint, France) over four cycles of 3 min lights on and 3 min lights off per cycle (epoch) for 24 minutes and a 2500 lux light intensity. After the 24-minute run for LPR, an acoustic startle response assay (900 ms of a 600 Hz frequency sound) was conducted using a ZebraBox for 2 minutes within the light phase on. Data was acquired using Viewpoint’s Zebralab software and analyzed in R using custom codes modified from previous work^1^. Dead and deformed embryos were excluded from the analysis. Phenotypes were observed and imaged under an Olympus TH4-100 equipped with a Lumenera Infinity 8-8 camera (Ottawa, Ontario, Canada). Phenotyping of embryos included assessments of each embryo's mortality, deformity, and developmental stages based on previous work^2^.

**2. Assessment of seizure like behavior**

The methods were adapted from a previous work^3,4^. Embryos were exposed to PFHxS from 2-120 hpf. Data collection focused on recording the frequency of specific behavioral events exhibited by fish in each experimental well over a 20-minute observation period. These events were categorized into three stages: Stage I, characterized by "Hyperlocomotion," indicating a dramatic increase in swim activity; Stage II, featuring "Whirlpool swimming," marked by rapid, whirlpool-like circling swim behavior; and Stage III, denoted as "Tonic seizure," involving brief clonus-like convulsions leading to loss of posture, such as the fish falling to one side and remaining immobile for 1-3 seconds. The scoring was conducted by an examiner blinded to the experimental conditions.

**3. Quantification of PFHxS Uptake in Zebrafish Embryos Using LC-MS Analysis Sample Collection- extended details.**

A total of 150 embryos exposed to PFHxS were collected at five days post-fertilization (dpf), with each experimental group consisting of n = 3 replicates.

**Sample Preparation**

Collected samples were rinsed twice with molecular-grade water to remove external contaminants.

**Homogenization**

Samples were homogenized by adding 1 ml of ultra-pure water (Wako) to the collected embryos and larvae. The homogenized samples were then transferred into 15 ml tubes for further processing.

**Addition of Sodium Hydroxide and Internal PFHxS Standard**

3.8 ml of 10 mM sodium hydroxide was added to each homogenized sample. An internal PFHxS standard of 20 ng/ml in methanol, 0.2 ml, was introduced to facilitate quantitative analysis.

**Sonication**

Sonication of the samples was performed at room temperature for 30 minutes to ensure thorough extraction of PFHxS and the internal standard.

**Centrifugation**

Following sonication, samples were centrifuged at 17,750 g for 10 minutes at 4°C to separate the particulate matter and cell debris from the supernatant. Each homogenized sample was divided into four equal aliquots to facilitate parallel processing. Each aliquot underwent centrifugation at 17,750 g for 10 minutes at 4°

**Sample Mixing**

Following centrifugation, supernatants from all four aliquots were carefully collected and combined into a single homogeneous sample in a 50 ml falcon tube and mixed with 9 ml of molecular grade water and vortexed to achieve homogeneity before analysis. Samples were frozen until further preparation.

**Solid Phase Extraction (SPE)**

Solid phase extraction (SPE) was performed according to the previous protocol from Ulhaq et al^5^ Briefly, Waters Oasis WAX 3 cc, 60 mg, 30 µm cartridges were preconditioned with 3 mL of 0.1% (m/v) ammonium hydroxide in methanol, then 3 mL of methanol, and then 3 mL of water. All solvents were Fisher Optima LC-MS grade. Thawed samples (14 mL) were loaded onto the cartridges and washed with 3 mL of 25 mM ammonium acetate buffer and then 3 mL of 50% aqueous methanol (v/v). PFHxS was eluted with 4 mL of 0.1% (m/v) ammonium hydroxide in methanol and dried down under nitrogen. Samples were reconstituted in 500 µL of 0.1% (m/v) ammonium hydroxide in methanol prior to LC-MS analysis.

**LC-MS Analysis**

Samples were injected (10 µL) in triplicate with an Agilent 1290 Infinity II UHPLC onto an Agilent ZORBAX Extend-C18 column (2.1 × 50 mm, 1.8 μm) maintained at 50 °C. The 10-minute gradient is displayed in the table below; Mobile Phase A was water (10 mM ammonium acetate) and Mobile Phase B was acetonitrile.

**Table S1.** LC gradient conditions.

| **Time (min)** | **MP A%** | **MP B%** |
| --- | --- | --- |
| 0.00 | 95 | 5 |
| 2.00 | 60 | 40 |
| 4.50 | 40 | 60 |
| 5.50 | 5 | 95 |
| 8.00 | 5 | 95 |
| 9.00 | 60 | 40 |
| 9.01 | 95 | 5 |

Following chromatographic analysis, samples were analyzed with an Agilent 6560 IM-QTOF operated in QTOF-only mode. Ionization was performed with an Agilent Jetstream (AJS) source operated in negative mode. Additional instrument conditions are included in the table below. PFHxS was detected as the [M-H]^-^ ion at *m/z* 398.937 with retention time of 3.20 mins. Extracted ion chromatograms (EIC) were generated for all samples and peak areas integrated and corrected for the included 20 ng/mL internal standard. A calibration curve ranging from 0.5-40 ng/mL was also generated to determine PFHxS in the individual samples.

**Table S2.** Instrumental parameters for the Agilent 6560 IM-QTOF.

| **Instrument Region** | **Instrumental Parameter** | **Experimental Value** |
| --- | --- | --- |
| **Source** | Gas Temp | 325 °C |
|  | Drying Gas | 12 L/min |
|  | Nebulizer | 20 psi |
|  | Sheath Gas Temp | 275 °C |
|  | Sheath Gas Flow | 10 L/min |
|  | VCap | 4000 V |
|  | Nozzle Voltage (Expt) | 1000 V |
|  | Fragmentor | 400 V |
|  | Oct 1 RF Vpp | 750 V |
| **High Pressure Funnel** | High Pressure Funnel Delta | -150 V |
|  | High Pressure Funnel RF | -150 V |
| **Trapping Funnel** | Trap Funnel Delta | -180 V |
|  | Trap Funnel RF | -150 V |
|  | Trap Funnel Exit | -10 V |
|  | Entrance Grid High | -96 V |
|  | Trap Entrance | -91 V |
|  | Trap Exit | -90 V |
|  | Trap Exit Grid 1 | -87.3 V |
|  | Trap Exit Grid 2 | -86.5 V |
| **Rear Funnel** | Rear Funnel Entrance | -240 V |
|  | Rear Funnel RF | -150 V |
|  | Rear Funnel Exit | -43 V |
|  | IM Hex Delta | 8 V |
|  | IM Hex RF | -600 V |
|  | IM Hex Entrance | -41 V |

**Table S3. Concentration of PFHXs within each sample (3 replicates). Pertains to Figure 1I.**

| Sample | Concentration (ng/mL) | Concentration (ng/embryo) |
| --- | --- | --- |
| DMSO 1 | 0 | 0 |
| DMSO 2 | 0 | 0 |
| DMSO 3 | 0 | 0 |
| Low 1 | 3.9 ± 0.4 |  |
| Low 2 | 4.8 ± 0.1 |  |
| Low 3 | 0.3 ± 0.1 |  |
| High 1 | 59.8 ± 7.8 |  |
| High 2 | 67.9 ± 4.3 |  |
| High 3 | 61.0 ± 3.3 |  |

**4. RNA seq**

RNA sequencing data generated by Novogene were analyzed using the iDEP.96 platform (<http://bioinformatics.sdstate.edu/idep96/> ). The read counts from RNA-Seq data were uploaded as a CSV file into the platform, and zebrafish (Danio rerio) was selected as the reference species. The variance stabilizing transformation (VST) option was applied to the Transform counts data for clustering & PCA during preprocessing. The default settings were retained for K-Means clustering and PCA. The DESeq2 method was utilized under the DEG1 section for differential expression analysis. The FDR cutoff was set to 0.05, with a minimum fold change threshold of 1.5. Factors and comparisons were specified according to the experimental design detailed in the uploaded file. A complete list of differentially expressed genes (DEGs), including FDR and fold change values, was downloaded for further analysis. A volcano plot was also generated to visualize the results in the DEG2 section. Pathway analysis was conducted using the KEGG pathway option within iDEP.96 under DEG2.

**5. Immunohistochemistry**

Immunohistochemistry was performed as previously described ^6^ on 16 micron retinal sections from treated 5 dpf larvae and their control siblings. Primary antibodies included mouse anti-Zpr-1 monoclonal antibody (1:200; University of Oregon Monoclonal Antibody Facility), rabbit anti-Rhodopsin antisera (1:5000, gift from David R. Hyde), and mouse anti-Proliferating Cell Nuclear Antigen (PCNA) monoclonal antibody (1:1000; clone PC10, Sigma-Aldrich, St. Louis, MO, USA). AlexaFluor goat anti-rabbit IgG 488 (1:500) and goat anti-mouse IgG 594 (1:500) were used as secondary antisera (Invitrogen-Molecular Probes, Eugene, OR, USA). Nuclei were labeled with TO-PRO-3 (Invitrogen-Molecular Probes, Eugene, OR, USA) at a 1:500 dilution. For TUNEL analysis of apoptosis, tissue sections were permeabilized ice-cold 0.1% NaCitrate/0.1% Triton X-100/1X PBS for 2 min and washed in 1X PBS for 5 min at room temperature (RT). Next, sections were incubated in 100 µL of labeling buffer (ApoAlert DNA fragmentation kit; Clontech International, Mountain View, CA, USA) for 10 min at RT, followed by humidified incubation with 50 µL of labeling mix (48 µL labeling buffer, 1 µL of 1 mM biotinylated dNTPs (New England Biolabs, Ipswich, MA, USA) and 1 µL of TdT enzyme (45 U/µL; ApoAlert DNA fragmentation kit; Clontech International, Mountain View, CA, USA), at 37 °C for 2 h. A 15 min RT wash with 150 µL of 2X saline-sodium citrate (SSC) was used to stop the reaction. Next, tissue sections were washed in 1X PBS and then incubated with Streptavidin conjugated to AlexaFluor 488 (1:200, Invitrogen-Molecular Probes, Eugene, OR, USA) and TO-PRO-3 (1:500, Invitrogen-Molecular Probes, Eugene, OR, USA) diluted with 1X PBS for 1 h in the dark. All tissue sections were mounted with Prolong Gold Antifade Reagent (Cat. no. P10144, Thermo Fisher Scientific, St. Louis, MO, USA).

**6. Pineal Gland Ablation:**

Zebrafish embryos were divided into four groups at 2 hours post-fertilization (hpf): a dimethyl sulfoxide (0.025 % DMSO) control group, a group treated with 20 mM metronidazole (Mtz) from 24-48 hpf, a co-exposure group treated with both PFHxS and 20 mM Mtz, and a PFHxS-only group. From 2-24 hpf, both the DMSO and 20 mM Mtz groups were maintained in DMSO, while the co-exposure and PFHxS groups were exposed to PFHxS.

.

At 24 hpf, all embryos were washed. The DMSO group remained in DMSO, while the 20 mM Metronidazole (Mtz) group (previously in DMSO) was transferred to 20 mM Mtz for pineal gland ablation, and the co-exposure group (previously in PFHxS) was moved to 20 mM Mtz for co-exposure from 24-48 hpf. The PFHxS group continued in PFHxS. At 48 hpf, a second wash was conducted, and the DMSO group remained in DMSO, the 20 mM Mtz group was returned to DMSO, the co-exposure group was transferred back to PFHxS, and the PFHxS group continued in PFHxS. The co-exposure with 20 mM Mtz was limited to 24-48 hpf for the purpose of pineal gland ablation.

At 120 hpf, phenotyping was performed, as well as recording for both LPR and LSR.

**References.**
